# Understanding the Mechanisms of *Salmonella* Typhimurium resistance to Cannabidiol

**DOI:** 10.1101/2023.04.27.538601

**Authors:** Iddrisu Ibrahim, Joseph Atia Ayariga, Junhuan Xu, Daniel A. Abugri, Robertson K. Boakai, Olufemi S. Ajayi

## Abstract

The emergence of multidrug resistance poses a huge risk to public health globally. Yet these recalcitrant pathogens continue to rise in incidence rate with resistance rates significantly outpacing the speed of antibiotic development. This therefore presents an aura of related health issues such as untreatable nosocomial infections arising from organ transplants, surgeries, as well as community acquired infections that are related to people with compromised immunity e.g., diabetic and HIV patients etc. There is a global effort to fight multidrug resistant pathogens spearheaded by the World Health Organization, thus calling for research into novel antimicrobials agents to fight multiple drug resistance. Previously, our laboratory demonstrated that Cannabidiol (CBD) was an effective antimicrobial against Salmonella Typhimurium (S. Typhimurium). However, we observed resistance development over time. To understand the mechanisms S. Typhimurium uses to develop resistance to Cannabidiol (CBD), we studied the abundance of bacteria lipopolysaccharide (LPS) and membrane sterols of both susceptible and resistant S. Typhimurium. Using real-time quantitative polymerase chain reaction (rt qPCR), we also analyzed the expression of selected genes known for aiding resistance development in S. Typhimurium. We discovered that there was a significantly higher expression of blaTEM, fimA, fimZ, and integrons in the CBD-resistant bacteria, and these were also accompanied by a shift in abundance in cell surface molecules such as lipopolysaccharide (LPS) and sterols.

## 1. Introduction

Antimicrobial resistance has successfully become a global health menace, and resistance are often acquired by bacteria through health-care-related incidence (HRI) orchestrated by multiple drug resistant (MDR) and extended drug-resistant pathogens (EDRP) such as *Klebsiella pneumonia, Staphylococcus aureus, Enterococcus faecium, Acinetobacter baumanii, Enterobacter spp*., and *Pseudomonas aeruginosa* [**1, 2, 3, 4**].

The recalcitrance of pathogenic bacteria indicates that millions of people are at risk of infection [**5, 6, 7**]. This has forced the Centre for Disease Control and Prevention to change their 2019 memorandum, from”a coming post-antibiotic epoch” to the fact that we are already in the era of stringent antibiotic resistance [**7**].

Understanding the mechanisms of antimicrobial resistance is a baffling task that challenges both scientific intellect and economics [**3, 8**], and several pharmaceutical companies have given up on antibiotic research. The perilous financial situation of antibiotic research and development means that there are fewer novel therapeutics underway, and not even a single novel antibiotic class has been developed and approved for clinical application since the 1960s, especially for Gram-negative bacteria [**3, 8, 9**].

*Salmonellosis* (an infection caused by *Salmonella*, often through contaminated food or water) is lethal and approximately 1.35 million infections, 26,500 hospitalizations, and 420 mortality cases are reported annually in the United States alone [**10, 11**]. Stomach cramps, diarrhea, and fever are some common symptoms of *salmonellosis*, and these symptoms develop based on the six-six principle (6 hours-6 days) which may last four to seven days in symptomatic people. Some people may be asymptomatic or may only show symptoms after several weeks of infection [**12, 13**]. *Salmonellosis* can be prevented when hygiene is strictly practiced such as eating foods that are pasteurized, regular hand washing, keeping kitchen and restrooms clean, etc. [**14, 15**]. However, like all other pathogens, *Salmonella* is capable of escaping hygienic protocols and developing resistance to most antibiotics [**16**].

To curtail the resistance of pathogenic bacteria, cannabidiol (CBD), the primary anti-psychoactive component of the hemp plant has increasingly been studied and employed. It is a small molecule with a molecular weight of 314 Da, consisting mainly of pentyl-substituted bis-phenol aromatic class [**17, 18**]. These aromatic classes are often linked to the alkyl-substituted cyclohexane terpene ring system [**19, 20**]. Among the dozens of cannabinoids extracted from the hemp plant, Cannabidiol (CBD) is one of them, and have proven to possess some bioactive components that act as antimicrobials [**21, 22**]. Though Cannabidiol (CBD) was first isolated in 1940, yet it was until 1963 that the full understanding of its structure was unveiled [**23, 24**]. Since the elucidation of its structure, Cannabidiol (CBD) has been rigorously tested and applied as an antidote to several diseases such as spasticity due to multiple sclerosis, depression, appetite stimulation, sleep disorders, glaucoma, psychosis, and anxiety disorder among many others. Another pharmacological relevance of Cannabidiol (CBD) is its potential application in neuroprotective and anti-inflammatory diseases [**25**]. Epidiolex® in the United States and Epidyolex in the European Union are so far the oil-based liquid formulations of Cannabidiol (CBD) that have been approved respectively by the US Food and Drug Administration in 2018 and by the European Medicines Agency in 2019. These formulations were critically evaluated and shown to play a role in the oral treatment of epilepsy disorders such as Lennex-Gustaut and Dravet syndrome [**26, 27**]. Cannabidiol (CBD) antimicrobial ability has been widely reported and shown to have minimum inhibitory concentrations (MICs) in the range of 1-5 μg/mL over Gram-positive bacteria, particularly *Streptococci* and *Staphylococci* and Gram-Negative bacteria such as *Salmonella* [**28**].

Notwithstanding, the rich applications of Cannabidiol (CBD) as an antimicrobial, most bacteria have developed strategies of resisting Cannabidiol (CBD) similar to most antimicrobials and this pose’s threat to public health [**29, 30**]. Recently we have demonstrated that Cannabidiol (CBD) has potent antimicrobial activity against two strains of *Salmonella (*Typhimurium and Newington), as compared to some common broad spectrum antibiotics such as polymyxin B, Azitromycin, and Kanamycin [**28, 31**]. However, we observed over time that *Salmonella* developed resistance against Cannabidiol (CBD). Given this, this present study seeks to elucidate the mechanisms that *S*. Typhimurium employs to develop resistance to Cannabidiol (CBD).

## 2. Materials and Methods

### 2.1 Media, chemicals, bacterial strains, and other reagents

All chemicals (Hexane, methanol –LC-MS (≥99.9%), water, and ethanol absolute proof (≥99.5%) used in this research were of HPLC grade. Dulbecco’s Modified Eagle Medium (DMEM) and Fetal Bovine Serum (FBS) were purchased from ATCC (Manassa, CO, USA). Our laboratory has demonstrated that *S*. Typhimurium is susceptible to CBD at micromolar concentrations (**28**). The two PCR ribotypes of *S*. Typhimurium used in this study were named CBD-resistant *S*. Typhimurium and CBD-Susceptible *S*. Typhimurium that were created from the lab-coded BV4012 strains. This BV4012 is *S*. Typhimurium LT2 strain MS1868 (a kind gift from Dr. Anthony R. Poteete, University of Massachusetts). The CBD-resistant strains were previously developed in our laboratory (**31**). To create CBD-resistant colonies, the *S*. Typhimurium strains were subjected to prolonged CBD pressure. The resistant colonies were picked and plated on LB agar premixed with CBD at 10 μg/ml final concentration. From hence all cultures of CBD-resistant strains received 10 μg/ml CBD concentration or more in their growth media. Luria Broth (BD, Difco, Franklin Lakes, NJ, USA) and Luria Agar (BD, Difco, Franklin Lakes, NJ, USA) were the media used for culturing and maintenance of the bacteria. The CBD extraction and purification process was carried out by Sustainable CBD LLC. (Salem, AL, USA) and has been described elsewhere (**28**). The Vero cells (ATCC, CCL 81) used in this study were obtained from BEI Resources (Manassas, VA, USA). The Vero cells were cultured and maintained at 37 °C, 5% CO_2_ in a T-25 cm^3^ flask using DMEM supplemented with 10% FBS and 1% penicillin-streptomycin-amphotericin (**32**).

### 2.2 Extraction of Lipids

The CBD-resistant strain has been previously developed in our lab [**31**]. CBD*-*resistant or CBD-susceptible *S*. Typhimurium cells were cultured to mid-log stage in LB broth only for the susceptible or CBD-tinted LB broth for the CBD-resistant strain, then 10 mL of the culture was then pelleted and resuspended with equal volume of the LB broth only, or LB broth tinted with CBD with final concentration of (0.01 mg/mL). The bacteria cultures were allowed to grow at room temperature for 6 h. The LB broth removed after centrifugation and the pellet washed with 1X PBS twice. The pellet was then resuspended in 5 mL 1X PBS and aliquoted into 2 mL Eppendorf tubes at 1 mL volume per tube. Bacterial lipids were extracted from the cells using the Bligh/Dyer procedure [**33**] with fewer modifications. Thus, 500 μL of chloroform:methanol (1:2 v/v) was added to the resuspended cells in the 2 mL Eppendorf tubes. The samples were then vortexed at high speed for 20 s each, then centrifuged at 2600 rpm for 5 min to pellet insoluble cellular debris. Then 500 μL of the resulting supernatant was decanted into new 2 mL Eppendorf tubes and 500 μL each of chloroform and PBS was added. The samples were vortexed for the second time for 10 s and centrifuged at room temperature for 5 min at 2600 rpm. The resulting two liquid phases were observed, then the organic lower phase was carefully transferred into a fresh 2 mL Eppendorf tube using sterile pipette tips. Then the upper aqueous phase was discarded.

### 2.3 Ergosterols quantification using UV-Vis spectrophometry

Sterols with conjugated double bonds, for example, ergosterol, have been demonstrated to have strong absorbances between 250 and 300 nm and have the capacity at this wavelength to detect at a limit of 6 ng of ergosterol. It has been demonstrated that ergosterols peaks at 295 nm and a shoulder at 265 nm (34). The amount of ergosterols in CBD treated *S*. Typhimurium samples or the untreated controls were quantified using the ABS Spectral Max UV-Vis spectrophometer. 500 μL of the extracted lipid was aliquoted into 1mL curvette tubes and the absorbance measured at 295 nm. Three replicate measurements were made for each treatment, and the absorbance averaged.

### 2.4 Mysristic, palmitic acid, palmitoleic acid, stearic acid, erucic acid, and Oleic acids quantification using UV-Vis spectrophometry

Mysristic, palmitic acid, palmitoleic acid, stearic acid, erucic acid were classified in this work as a single group of sterols since they all have their absorbance at 208 nm (**35**). Thus this group of sterols quantification were made via measuring the absorbance of the extracted lipids at 208 nm. Oleic acids however are known to have absorbance at 330 nm (**36**), thus, the quantification of oleic acid was carried out via measuring absorbance at 330 nm.

### 2.5 Unsaturated and other uncatergorized bacterial membrane sterols quantification using UV-Vis spectrophometry

#### 2.5.1 LPS Extraction protocol

To compare the relative quantities of unsaturated and saturated bacterial membrane sterols, membrane unsaturated fatty acids were extracted using the Lipid Assay Kit (ab242305 Abcam, USA), the lipids were aliquoted into 1 mL cuvette tubes and were measured at an absorbance of 540 nm using the UV-Vis spectrophometer. For all other uncategorized membrane sterols, absorbance at 314 nm was used to quantify them.

### 2.6 Quantitative PCR Analysis of Gene Expression

To investigate the effect of CBD on the antibiotic resistance genes in *S*. Typhimurium, the total RNA of CBD-resistant and CBD-susceptible *S*. Typhimurium strain were (*n* = 3) were extracted using the Rneasy kit (Qiagen Sciences, Maryland, USA), after an overnight culture in CBD tinted LB broth (for CBD-resistant strain) and LB broth only (for CBD-Susceptible strains. After extraction, the RNA was dissolved in RNase-free water, and the concentration was determined by Nanodrop (1000 spectrophotometer, ThermoFisher Scientific, Madison, WI, USA). Following the RNA extraction, 100 ng of RNA from each sample was then reverse transcribed into cDNA using SSIV Cell Direct cDNA Synthesis kit (lot# 00916637, Invitrogen, ThermoFisher Scientific, Vilnius, Lithuania). Subsequently, the cDNA was subjected to quantitative PCR analysis for the following antibiotic markers; int1, int2, int3, blaTEM, fimA, fimZ, STM0959, STM0716 and the spy genes, using CFX96™ real-time PCR (Bio-Rad Laboratories, Hercules, CA, USA) following similar procedures as in our previous work (**37**).

The primers for int1, int2, int3, blaTEM, fimA, fimZ, STM0959, STM0716 and the spy genes were designed and ordered from ThermoFisher Scientific, and the DNA gyrase B (gyrB) gene served as the housekeeping gene. Real-time QPCR analysis of int1, int2, int3, blaTEM, fimA, fimZ, STM0959, STM0716 and the spy genes and the *S*. Typhimurium housekeeping gene (gyrB) was carried out using the primers listed in Table 1 and powerup™ SYBR™ Green Master Mix (lot# 00914521). Comparisons of target gene expression in the samples were carried out using the cycle threshold (Ct) normalized to the housekeeping gene gyrB.

**Table 1:**
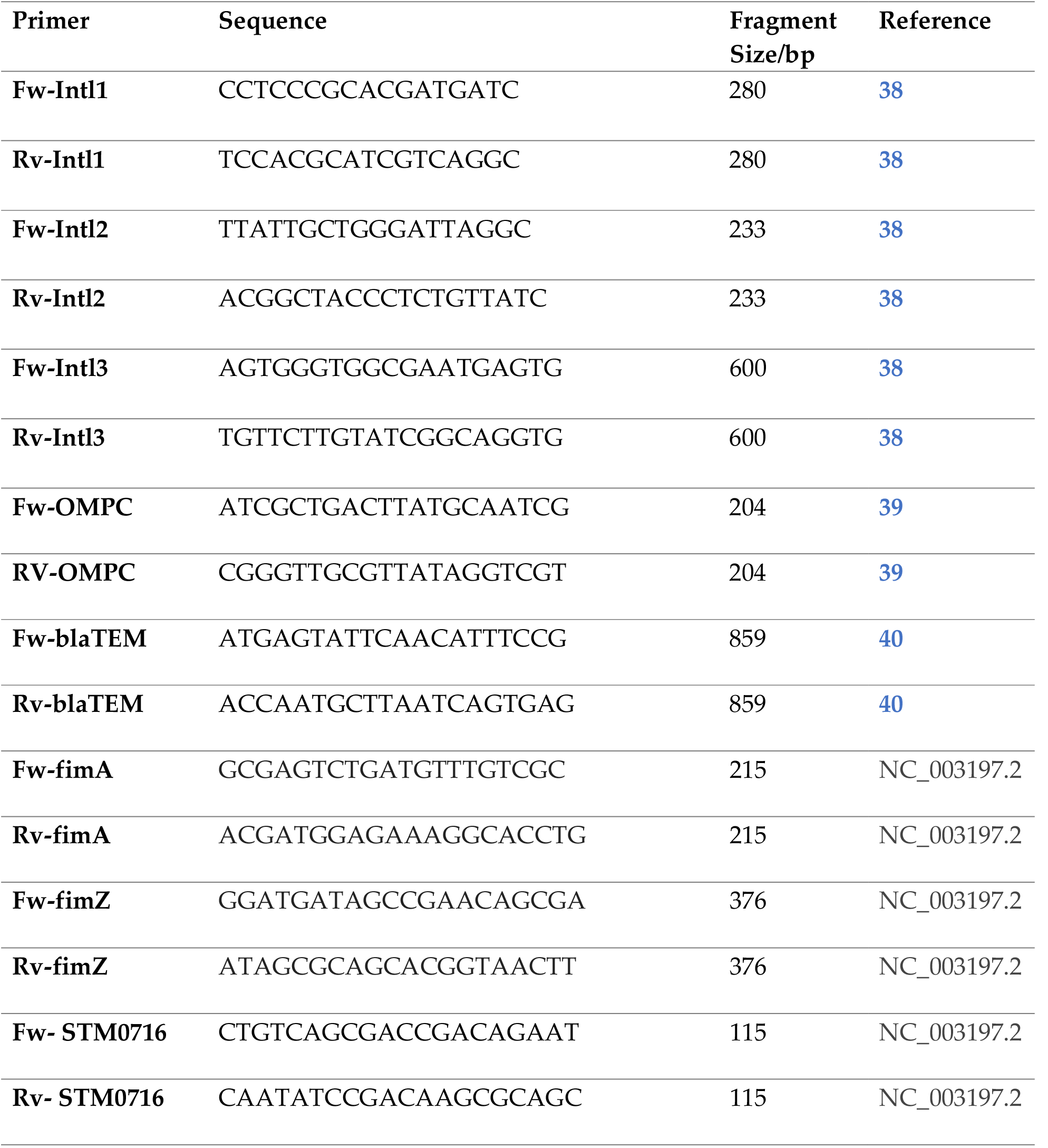
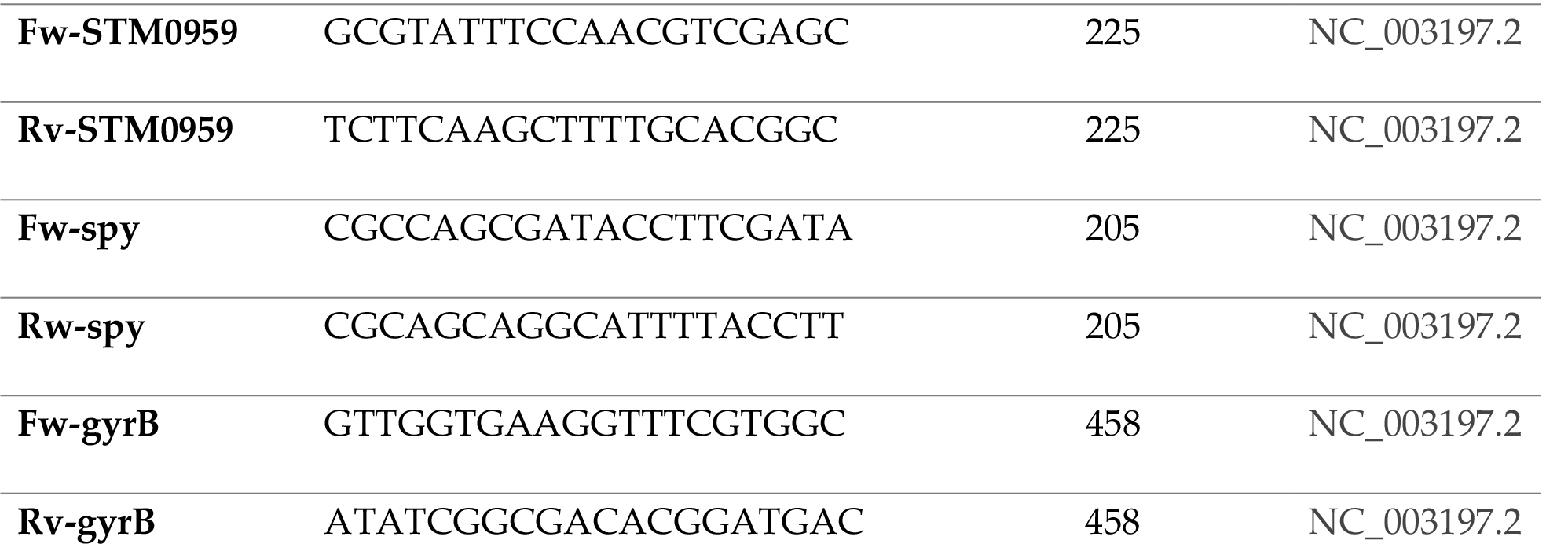
List of the primers used in the PCR analysis of int1, int2, int3, blaTEM, fimA, fimZ, STM0959, STM0716 and the spy genes expression in CBD-resistant and CBD-susceptible strain of *S*. Typhimurium.

### 2.7 Anti-invasion Assay

To assess the ability of CBD-resistant *S*. Typhimurium to resist the CBD as a prophylactic agent in Vero cell lines, Vero cells at a density of 1 × 10^5^ were cultured at 37°C in an atmosphere containing 5% CO_2_ in DMEM media supplemented with 10% FBS (Gibco, Grand Island, NY, USA) in 96-tissue culture plate (Corning Costar, Milano, Italy) for 24 h. Then the media removed, and 300 μL of CBD-resistant or CBD-susceptible *S*. Typhimurium cells at an OD_600_ of 0.3 were added to the growing Vero cells. This was then immediately followed with the administration of CBD at 100 μg/mL, or media control supplemented DMEM media only. Infection was allowed for 30 min following a similar procedure by Ayariga et al., and Manini et al., [**41, 42**]. Final concentrations of CBD in the culture media were 50 μg/mL in all the experimental group, with the control receiving the DMEM media placebo. Before the infection of the Vero cells, *S*. Typhimurium strain (both CBD-resistant and CBD-susceptible) were harvested after 6 h growth and pelleted via centrifugation at their mid log phase. Bacteria pellets were washed thrice with 1X PBS and resuspended in DMEM media to an OD_600_ of 0.3. To investigate the capacity of the CBD-resistant *S*. typhimurium to resist the killing ability of CBD and thus survive and invade the monolayer Vero cells, the wells containing the infected Vero cells, or the controls were washed 3 times with 1X PBS after the 30 min treatment with CBD, and then stained with acridine red. Bacterial cells stains red in the presence of acridine. Thus, bacterial cells that were able to resist the killing ability of the CBD treatment and hence invaded the Vero cells were captured whereas, *S*. Typhimurium which were susceptible to CBD died and could not invade the Vero cells were washed and thus could not be captured.

### 2.8 LPS Extraction

Gram-negative bacteria are known for their Lipopolysaccharides(LPS) that form a major component of their outer membrane. LPS is critical in resistance to antibiotics, phagocytosis, serum, and serves as an outer membrane permeability barrier. It serves as a major receptor for majority of phages, e.g. the well-known Salmonella P22 and Epsilon 34 phages (30). In this experiment we extract *S*. Typhimurium LPS using the LPS extraction kit (iNtRON Biotechnology, Inc.), following the manufacturers protocol. In brief, S. Typhimurium cells grown in CBD tinted LB broth (for CBD-resistant strain) or LB broth only (for CBD-Susceptible strain) to mid-log phase were centrifuged at 13,000 rpm for 1 min to pellet cells. The bacterial pellet was then washed thrice with 1X PBS to remove all traces of media. 1 ml of the bacterial Lysis Buffer was added to the pellet, and the pellet loosened and vigorously vortexed. This was then followed by the addition of 200 ml of chloroform, and then vortexed again for 20 sec. Then, sample was incubated at for 5 min at 25 °C. Following incubation, the sample was then centrifuged at 13,000 rpm for 10 min at 4 °C, and 400 ml of the supernatant was carefully transferred into a fresh 1.5 ml tube. To the 400 ml of the supernatant, 800 ml of the purification buffer was added and mixed thoroughly, then incubated for 10 min at -20 °C in the freezer. This was then followed by centrifugation at 13,000 rpm for 15 min at 4 °C. The upper layer was then carefully discarded to obtain LPS pellet. Then, 1 ml of 70% EtOH was added to wash the LPS pellet by inverting the tube 3 times. Then the mixture was centrifuged for 3 min at 13,000 rpm at 4°C and the supernatant discarded to obtain the washed LPS pellet. The LPS pellet was then resuspended in 50 ml of 10mM Tris-HCl buffer(pH 8.0) and dissolved thoroughly by boiling for 2 min. This was then followed by treatment with 75 μg/ml concentration of proteinase K for 50 °C for 30 min.

### 2.9 Statistical Analysis

Statistical analyses were performed using OriginPro Plus version 2021b (OriginLab Corporation, Northampton, MA, USA) or Microsoft Excel 2013 (Microsoft, Redmond, WA, USA) and statistical significance tested via the paired t-test. Immunofluorescence images were analyzed and image qualities readjusted by image J (NIH free open-source software). All statistical data were expressed as mean ±Standard error of mean.

## 3. Results

### 3.1 Comparisons of lipopolysaccharide, ergosterols, mysristic palmitic and Oleic acids of Susceptible and Resistant strain of S. Typhimurium

Quantification of the target liposaccharides (LPS) shows significantly higher abundance of the liposaccharides (**Figure 1A**) in the CBD-resistant strain as compared to the CBD-susceptible strain. Also, comparative abundance of the lipids such as ergosterols (**Figure 1B**), mysristic palmitic (**Figure 1C**) and oleic acids (**Figure 1D**) depicts higher abundance in the resistant strain of *S*. Typhimurium as compared to the susceptible strain (**Figure 1A-D)**.

**Figure 1.**
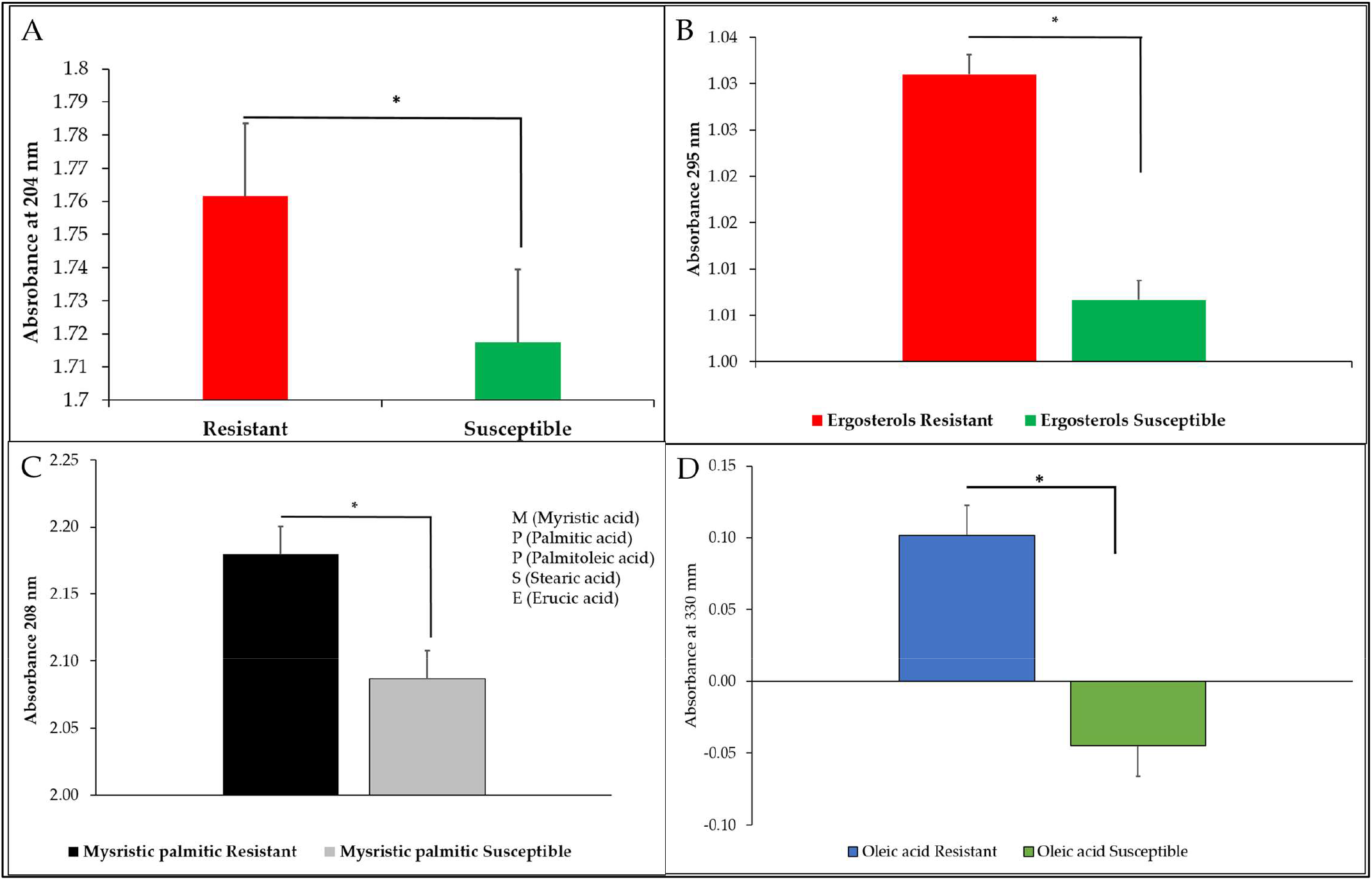
Relative abundance of A) LPS, B) Ergosterols C) Mysristic palmitic acid D) Oleic acid resistance of susceptible and resistant *S*. Typhimurium. Ergosterols p-value = 0.389263, Palmitic p-value = 0.001483, Oleic acid p-value = 0.001687, Unsaturated FA p-value = 0.009726, Other membrane sterols p-value = 0.002726

### 3.2 Membrane fatty acids Composition of Susceptible and Resistant S. Typhimurium

A pie chart showing the membrane sterols compositions of both resistant and susceptible strain of *S*. Typhimurium. The chart shows that the unsaturated fatty acids were highly abundant in both the resistant and susceptible strain representing 23% and 22% respectively. The next most abundant membrane fatty acid is the mysristic palmitic acids in both the resistant and susceptible strain representing 28% and 17% respectively. With the least abundant membrane fatty acid being oleic acids representing 1% in the resistant and 0% in the susceptible strain of *S*. Typhimurium (**Figure 2**).

**Figure 2.**
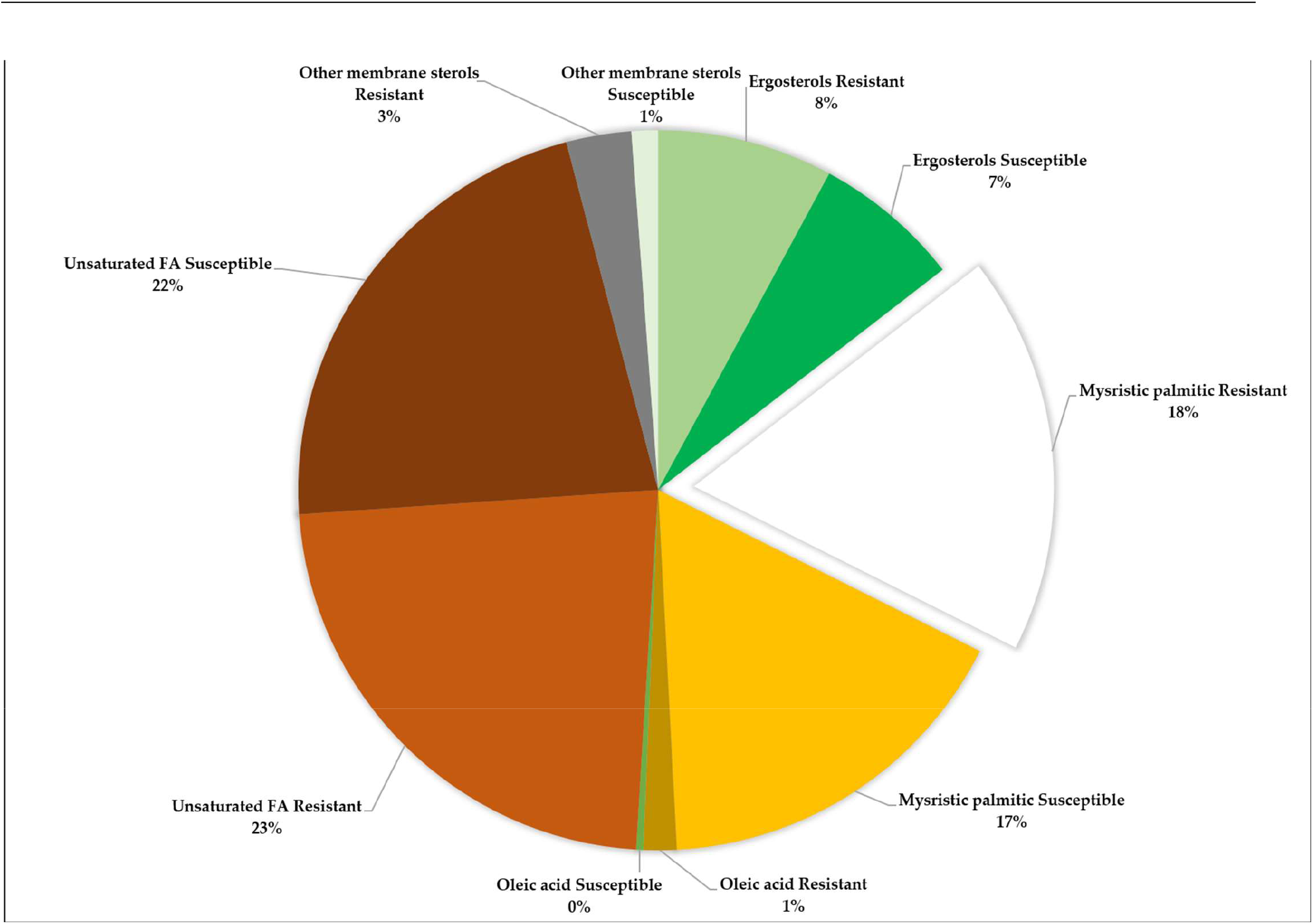
Relative abundance of CBD-Susceptible and CBD-Resistant *S*. Typhimurium.

### 3.3 *Immunofluorescence panels of S*. Typhimurium *infection of Vero cells and treatment with CBD*

Immunofluorescence imaging of both CBD-resistant and CBD-susceptible *S*. Typhimurium strain treated to CBD. **Figure 3A** shows the bacteria only stained with acridine red dye with gives the bacteria a red fluorescence under Texas Red filter. The immunofluorescence studies revealed that susceptible strain of *S*. Typhimurium were killed by the CBD treatment as shown in **Figure 3B** lane 2 (at Texas Red) whereas, the resistant strain of *S*. Tphimurium in lane 3 remained alive, thus showing high red flourescence, hence, demonstrating the presence of intracellular *S*. Typhimurium in the Vero cells.

**Figure 3.**
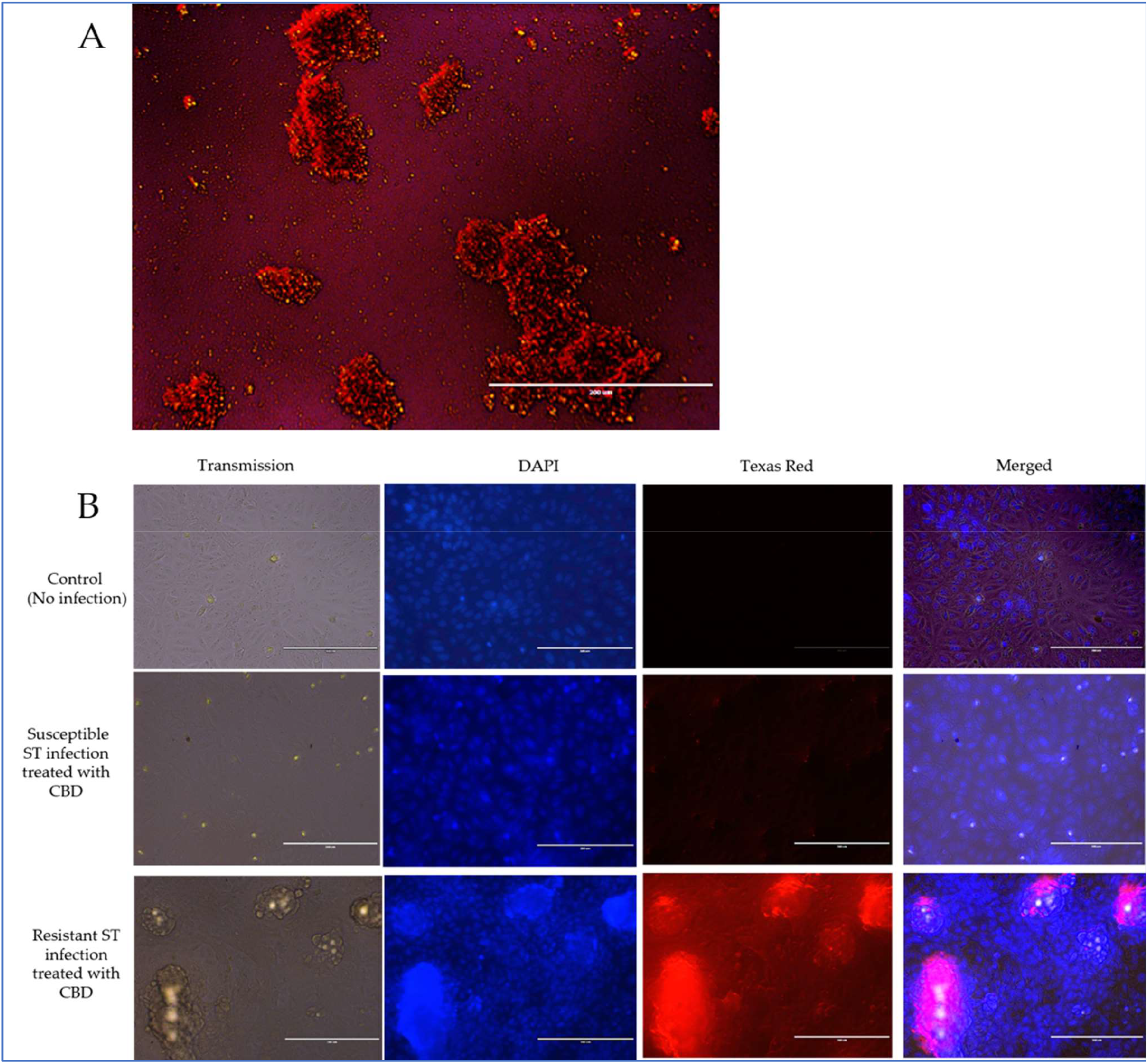
Panels showing the Immunofluorescence images of *S*. Typhimurium infection of Vero cells and counter treatment with CBD.

### 3.4 Gene expressions of susceptible and resistant strain of S. Typhimurium

To understand the underlying possible genetic drivers that caused the CBD resistance, we investigated some common antibiotic resistance genes such as, int2, int3, blaTEM, fimA, fimZ, STM0959, STM0716 and the spy genes expression in CBD-resistant and CBD-susceptible strain of *S*. Typhimurium shown in **Figure 4A-C**. Comparing the integrons (Int), Int1 and Int3 were upregulated in the susceptible strain, whiles Int2 showed high expression in the CBD-resistant strain as against the resistant strain of the *S*. Typhimurium (**Figure 4A-C)**.

**Figure 4.**
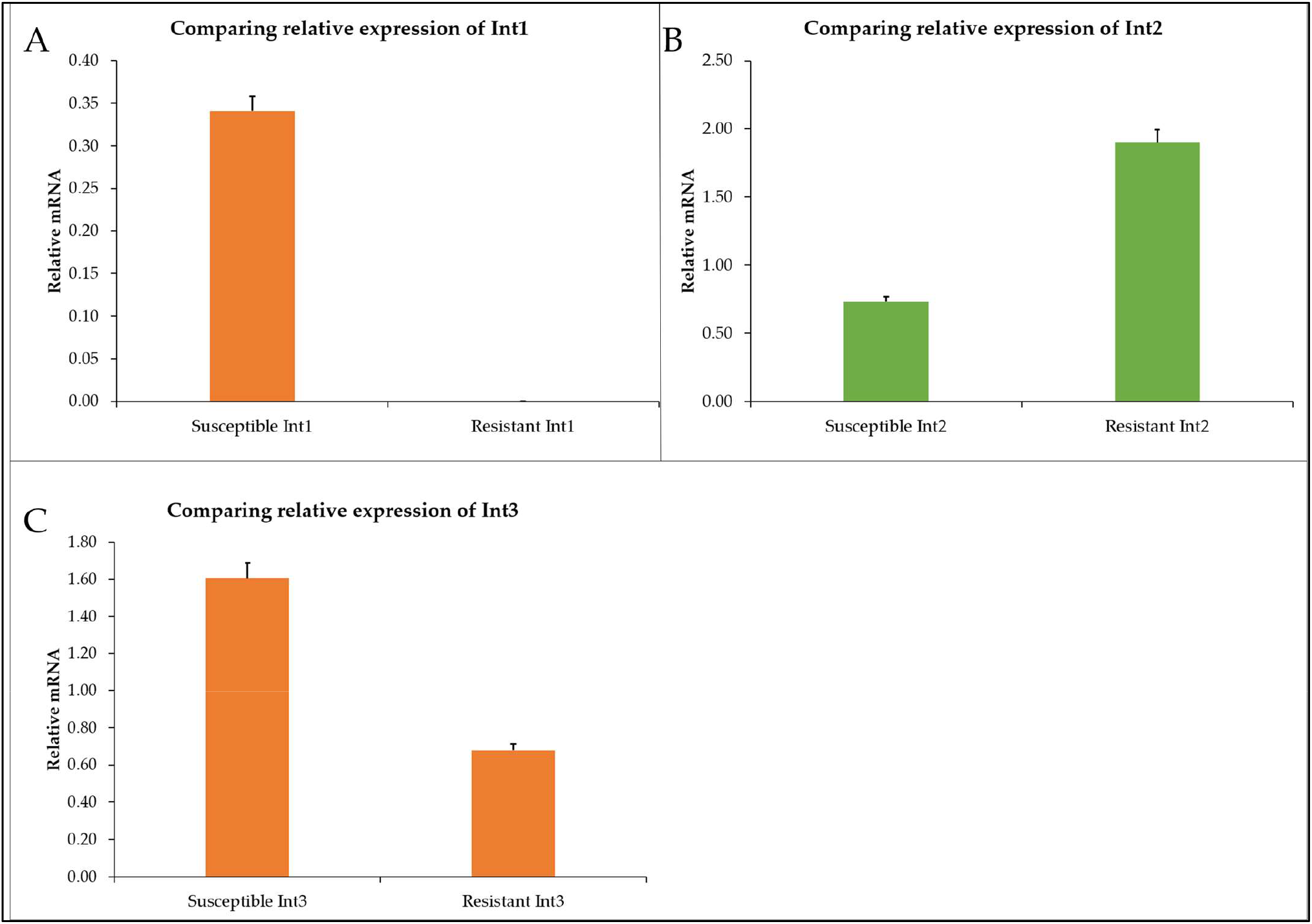
Relative mRNA expression of Integrons in susceptible and resistant *S*. Typhimurium A) Integron 1 B) Integron 2 C) Integron 3.

Similarly, genes coding for bacteria fimbriae (fimA and fimZ) showed higher expression in the CBD-resistant strain of *S*. Typhimurium as compared to the CBD-susceptible strain (Figure **5A-B**).

**Figure 5.**
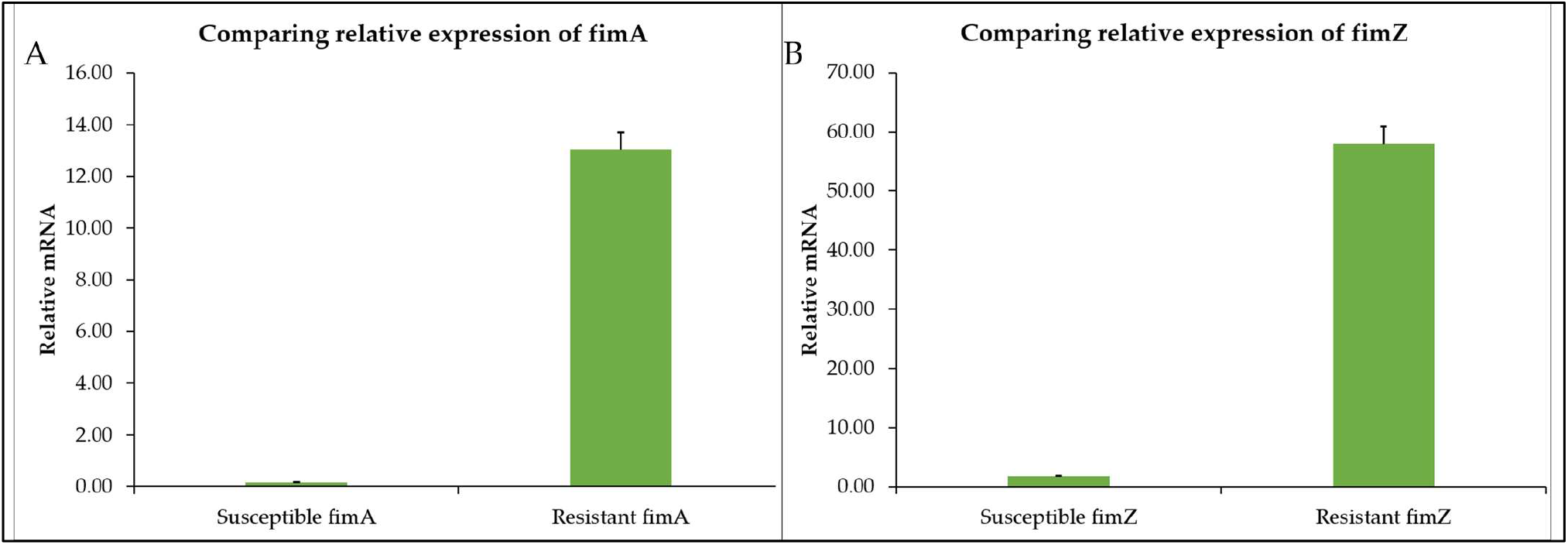
The relative mRNA expression of A) fimA, B) fimZ in susceptible and resistant *S*. Typhimurium.

Leucine-response (Lrp) transcriptional regulator with entry number (STM0959) were upregulated in the CBD-resistant strain and downregulated in the CBD-susceptible strain (**Figure 6A)**. Another gene of interest was the DNA recombinase (KEGG Entry number: STM0716) were downregulated in the CBD-resistant strain, however, in the case of the CBD-susceptible strain this gene was highly expressed (**Figure 6B)**.

**Figure 6.**
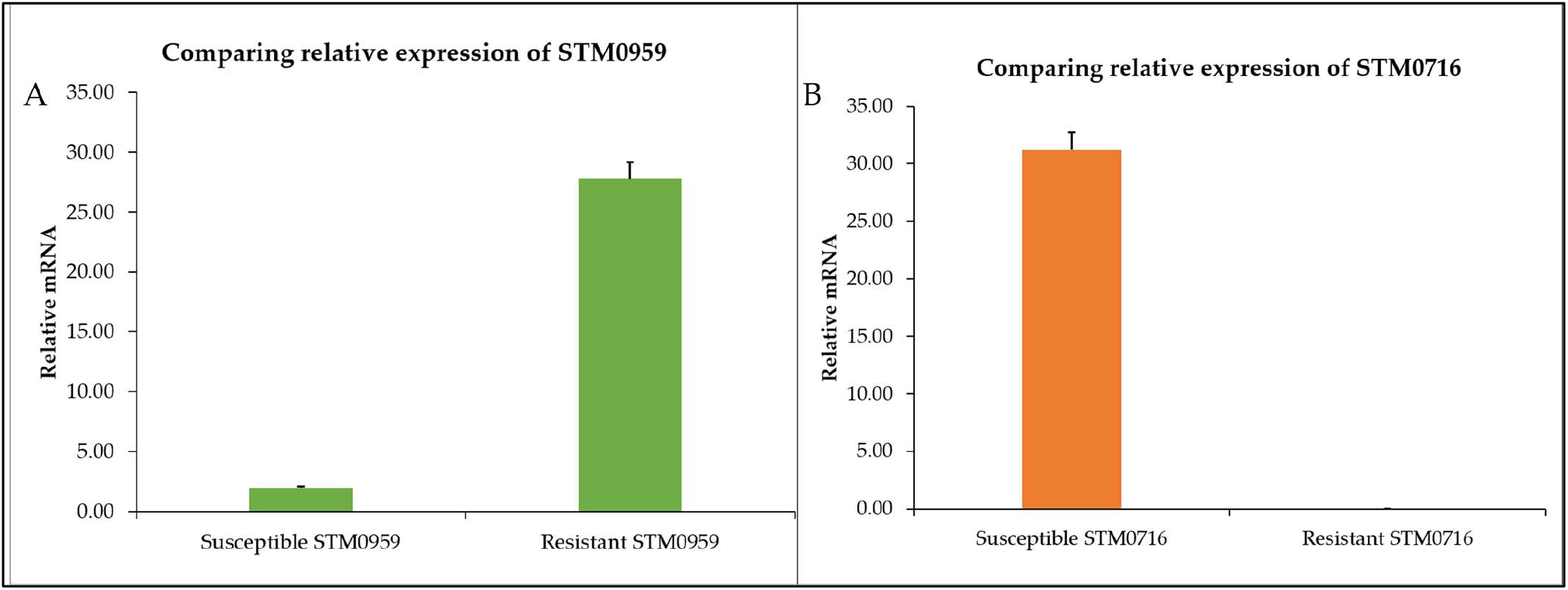
Relative expression of A). STM0959 and B). STM0716 in CBD-susceptible and CBD-resistant *S*. Typhimurium.

Furthermore, comparing beta-lactamase (blaTEM) gene expression in both strain shows that the gene was highly expressed in the CBD-resistant strain as compared to the CBD-susceptible strain of *S*. Typhimurium (**Figure 7A**). Close to 200 relative mRNA expression levels is noted in the blaTEM expression in the CBD-resistant strain, whereas negligible expression levels of blaTEM was recorded for CBD-susceptible strain. Similarly, outer membrane protein C (ompC) gene expression levels of CBD-resistant strains was depicted to be approximately twice the expression levels of the CBD-susceptible strain (**Figure 7B)**. In the case of the spy gene, it showed higher expression in the CBD-resistant strain than in the CBD-susceptible strain of *S*. Typhimurium (**Figure 7C**).

**Figure 7.**
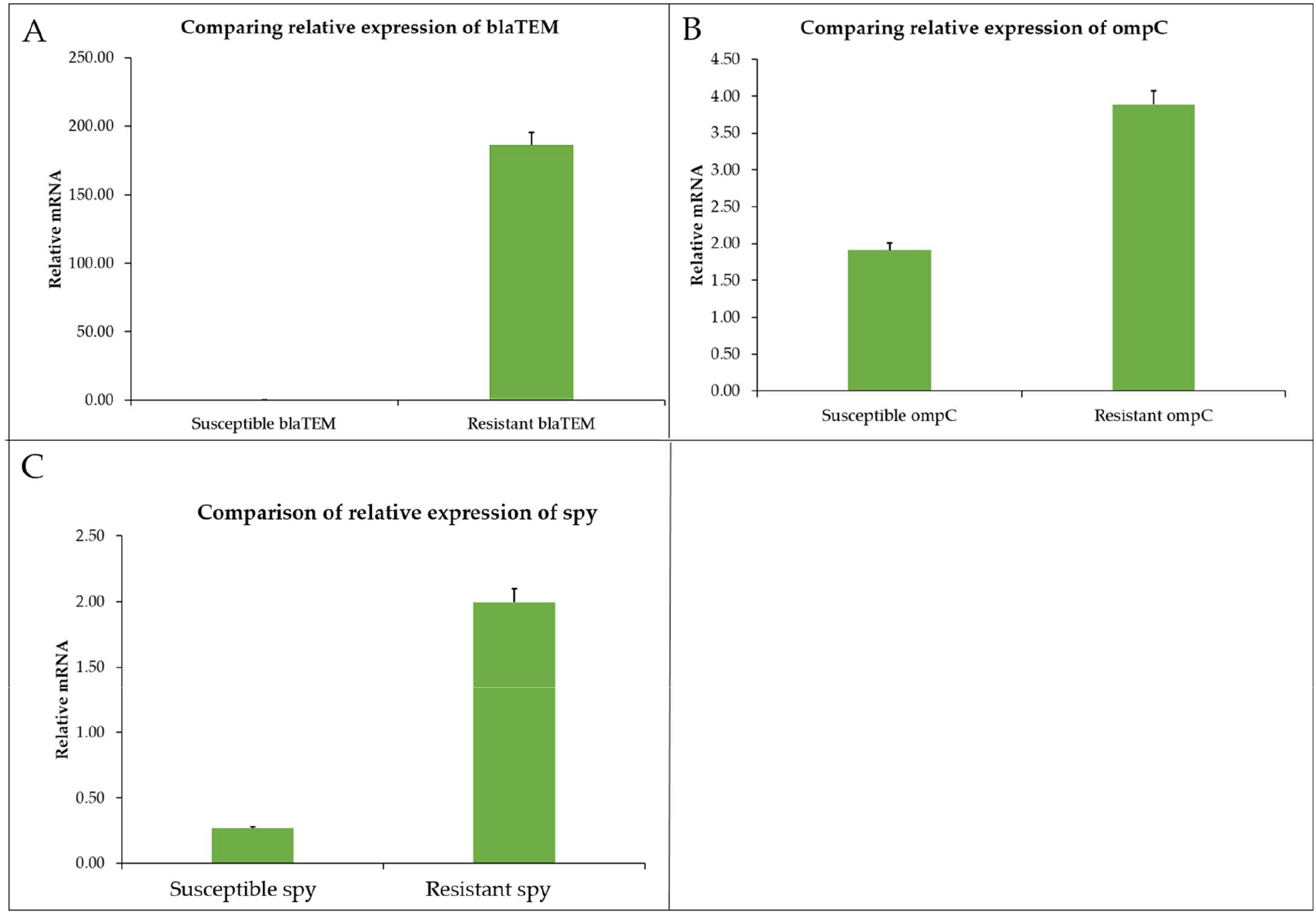
Relative expression of A) blaTEM, B) ompC, C) spy in CBD-susceptible and CBD-resistant *S*. Typhimurium.

The pie chart below represents a holistic view of all the selected genes considered in this study. The chart shows beta-lactase (blaTEM) which occupied more than half of the pie chart was the highest expressed gene in the CBD-resistant strain of *S*. Typhimurium as compared to the other genes. The second highest expression is recorded for fimZ in the CBD-resistant strain, followed by STMO716 in the CBD-susceptible strain, STM0959 and fimA in the CBD-resistant strain which also shared significantly high expression profiles (**Figure 8**).

**Figure 8.**
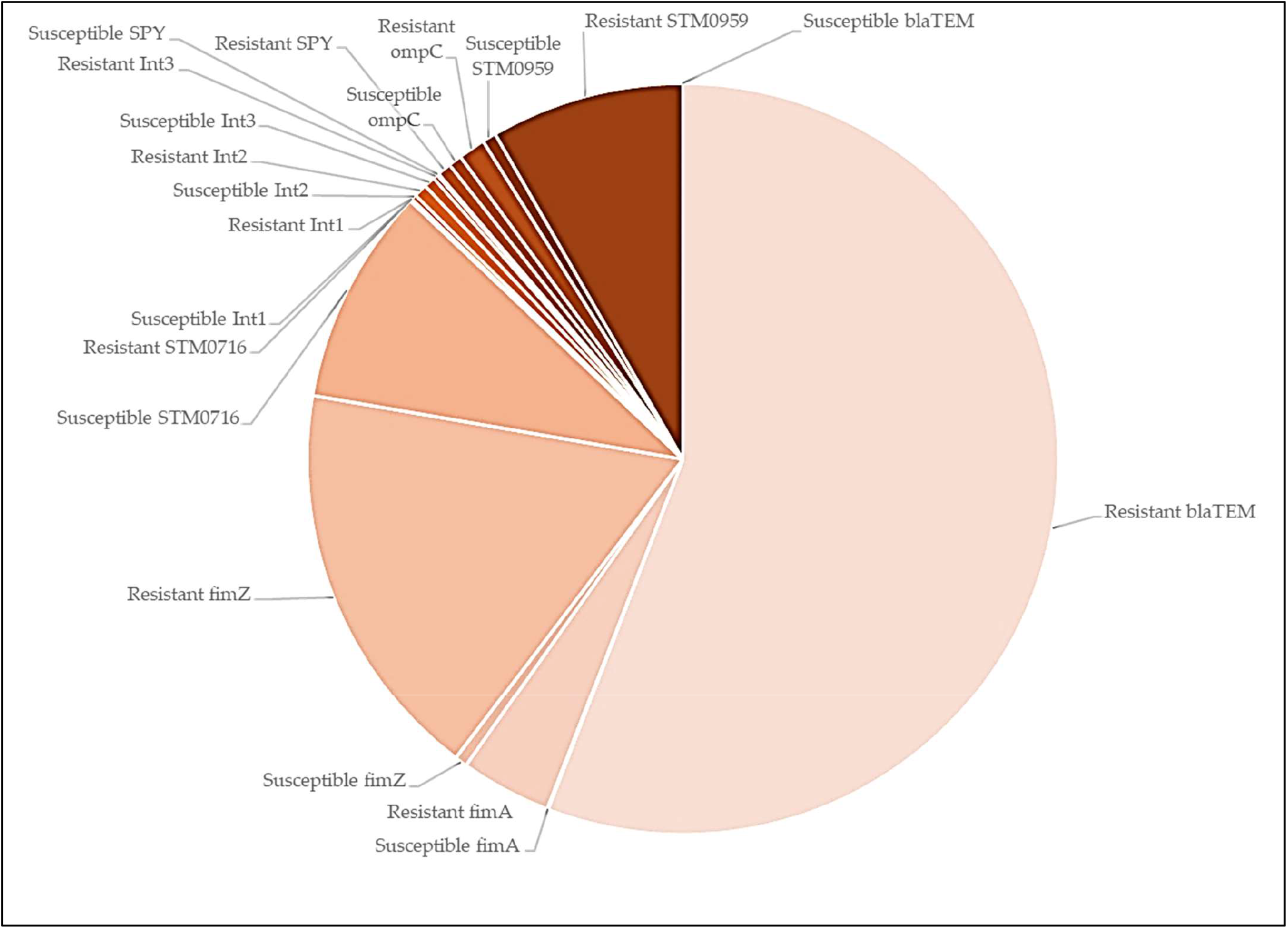
Pie chart of all genes under study that are expressed in CBD-susceptible and CBD-resistant strains of *S*. Typhimurium.

### 3.5 *Interactions of S*. Typhimurium *resistance genes with closely related genes*

To understand in details other genes implicated in *S*. Typhimurium resistance development, we carried out predictive studies using string network analysis. We determined that other genes closely interact with the selected set of genes under investigation. We found from the prediction analysis that fimA is closely interacting with fimH, fimD, fimF, fimC and fimI (**Figure 9A**). The spy gene is closely interacting with cpxA, cpxR, mdtA, STM2535, dsbA, nlpB and ybaJ genes (**Figure 9B**). ompC is closely interacting with yaeT, bluB, ompA, ompX, ompR, and osoxS genes (**Figure 9C**). And fimZ is closely interacting with fimC, fimF, fimh, STM3012, torS, rcsC, umY, and ssrB genes (**Figure 9D**).

**Figure 9.**
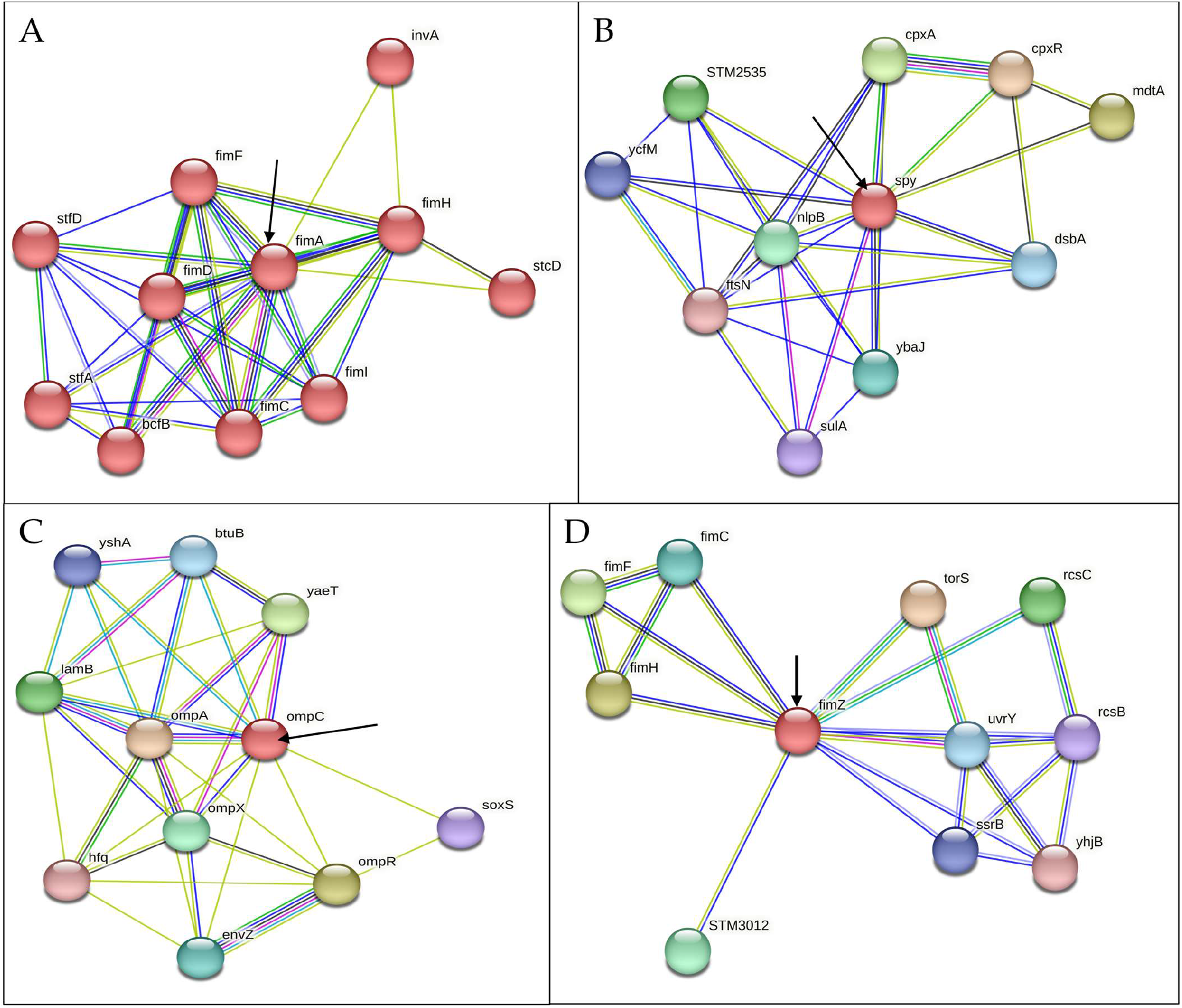
String-network depicting Interactions of selected CBD-resistance genes with other closely related genes A) fimA, B) spy, C) ompC, D) fimZ.

## 4. Discussion

Bacteria genetic plasticity has made them remarkable and permits them to adjust to varied environmental stresses imposed on them including antibiotics that may be deleterious to their existence [**43**]. Bacteria typically acquire certain intrinsic resistance mechanisms through antimicrobial-producing microorganisms that they share the same ecological niche with, which enables them to withstand antibiotic stresses [**44**].

In our previous studies, we demonstrated that repeated exposure of *S*. Typhimurium to CBD resulted in CBD-resistance development [**31**]. In this study, our primary objective was to ascertain the mechanisms *S*. Typhimurium employs to develop resistance to the potent CBD. We discovered that *S*. Typhimurium CBD-resistance might have arisen from a variety of factors such as LPS and membrane sterols modification and higher expression of specific genetic factors including blaTEM, fimA, fimZ, and lrp (SMT0959) genes.

Lipopolysaccharide (LPS) is the main structural component of the outer membrane and cell envelop of most Gram-Negative bacteria, which is clinically essential due to its role in bacteria pathogenicity and infection [**45**]. We have observed in this study that there is a distinct difference in the LPS abundance of the CBD-resistant *S*. Typhimurium strain as compared to the CBD-susceptible strain (**Figure 1A**). In this study we recorded higher abundance of LPS in the CBD-resistant strain. Abundance of the LPS enhances the outer membrane as a useful permeability barrier for small molecules, as well as hydrophobic molecules that could have crossed the phospholipid bilayers. This property of LPS modification according to Bertani and Ruiz causes significant resistance development of most Gram-Negative bacteria to several antimicrobial agents [**45**]. Cytoplasmic membranes are the defining membrane that separates cells’ internal components from the extracellular matrix and plays a crucial role in the transduction of energy and the conservation of solute transport, which regulates cell metabolism [**46**]. Membranes also function as stress sensors and are capable of transducing signals that might affect genes responsible for defense [**46**]. Comparing the membrane fatty acids of the CBD-resistant and CBD-susceptible *S*. Typhimurium strains reveals some significant and observable differences in unsaturated fatty acids, ergosterols, mysristic and palmitic acid, oleic acids, and other membrane sterols as shown in **Figure 2**. These differences imply that membrane fatty acids play an enormous role in bacteria adaptation to CBD. Similar results have been obtained by Ayari et al., [**46**] when they subjected *Bacillus cereus* and *Salmonella typhi* to gamma irradiation treatment to measure the alterations in the membrane fatty acid composition of the two bacteria species. Additionally, outer membrane proteins (OMP) protect Gram-Negative bacteria from deleterious environments [**46**]. Simultaneously, several proteins, solutes and other signal transductions receptors are critical components of the outer membrane, which is critical to the survivability of bacteria cells. OMP acts as permeability barriers for the exchange of nutrients, passage of toxins, and antibiotics [**47**]. OmpC in particular is associated with multidrug resistance [**48**]. **Figure 7B** shows that the ompC gene was highly expressed in the CBD-resistance *S. typhimurium* strain compared to the CBD-susceptible strain. A study conducted by Liu et al. [**49**] to understand the role ompC plays in resistance development in *E. coli* by mutating *E. coli* and subsequently treating it with carbapenems and cafepime observed that mutated *E. coli* showed decreased susceptibility to carbapenems and cafepime. In our study, ompC also interacts with other closely related OMPs such as ompA, ompX, and ompR, which are known to be anchor proteins, and plays a structural role by maintaining the integrity of the bacteria cell surface. These proteins especially ompA provide cells physical connection between underlying peptidoglycan layers and cell outer membranes (**Figure 9C**). ompC also interacts with another crucial outer membrane protein such as ompX, which is part of a family of proteins known to be highly virulent and can destabilize the defense mechanism of its host [**50**]. These related proteins are crucial for the survival of *Salmonella* in macrophages (**Figure 9C**) [**50**]. While many pathogens devise new methods of invading host cells, bacteria adhesions serve a crucial function in maximizing viability and survival options. This is achieved by employing type 1 fimbriae, which are markedly varied. *S*. Typhimurium produces type 1 fimbriae, which gives them a firm grip to a diverse array of cells and confer them the capacity to withstand stressors [**51**]. In this study, we considered the expression of the fim genes (fimA and fimZ) to elucidate their role in CBD-resistance development. Both fimA and fimZ were highly expressed in the CBD-resistant strain and were almost absent in the CBD-susceptible strain (**Figure 5**). Higher fim expression allows the bacteria firm attachment and grant it resistibility. Althouse et al., [**51**] studied the type 1 fimbriae in *S*. Typhimurium and concluded that type 1 fimbriae were particularly responsible for *Salmonella* attachment to enterocytes and promoted intestinal colonization. They however did not observe any association between type 1 fimbriae and intracellular survival of the *S*. Typhimurium. Avalos et al., [**52**] also studied how type 1 fimbriae help *E. coli* to evade extracellular antibiotics. They found that type 1 fimbriae increased *E. coli* survival chances in macrophages. Additionally, after exposing *E. coli* to gentamicin, they recorded those type 1 fimbriae increased internalization and adhesion to macrophage. The fims are the structural component (main subunit) of the fimbriae. fimA and fimI are considered as adhesive or regulatory genes, and fimF is an adapter gene. In this study fimA was highly expressed (**Figure 9A**). Zeiner et al., [**53**] assessed fimA, fimF, and fimH to see if they were necessary for assembling type 1 fimbriae in *S*. Typhimurium. They achieved this by mutating the fim (A, F, and H) genes and examining their capacity to produce surface-assembled fimbriae. Interestingly, they discovered that *S*. Typhimurium mutants were unable to assemble fimbriae, which implies that these genes are required to produce fimbriae in *S*. Typhimurium. fimZ together with other related fim genes such as fimY activates fimA proteins all play crucial roles in the fimbriae structure assembly (**Figure 9D**). Avalos et al., [**52**] investigated the role of fimW, fimY, and fimZ in modulating the expression of type 1 fimbriae in *S*. Typhimurium. They indicated that fimZ presence could enhance the expression of hilE, which is reported to be the repressor of SP11 gene expression.

Integrons are genetic elements that bacteria employ to withstand and evolve rapidly from stressors by the expression of new genes. The expressed genes are incorporated into a site-specific genetic structure known as gene cassettes that usually carries a non-promoter open reading frame combined with a recombination site. Our study explores various integron genes (Int1, Int2, and Int3) to elucidate their role in supporting *S*. Typhimurium CBD-resistance development (**Figure 4**). Only Int2 was shown to be highly expressed in the resistant strain, while both Int1 and Int3 were lowly expressed in the CBD-resistant strain as compared to the CBD-susceptible strain. Deng et al., [**54**], critically reviewed the role of integrons (class 1, 2, and 3) in antibiotic resistance by pathogenic bacteria, and substantiate that integrons do indeed play a role in resistance development. Jones-Dias et al., [**55**] studied the architecture of class 1, 2, and 3 integrons from Gram-Negative Bacteria (*Morganella morganii, E coli*, and *Klebsiella pneumoniae*) recovered from fruits and vegetables. They stated that the diverse integrons constituents were associated with transposable elements and led to the identification of varied integron promoters such as PcW, PcS, PcH1, and PcWTNG-10. Furthermore, Firoozeh *et al*., [**56**] used molecular techniques to characterize class 1, 2, and 3 integrons in clinical multidrug resistant *Klebsiella pneumonia* isolates. They studied the antibiotic resistance patterns of the isolates and found that about 150 isolates carried the Int1 gene, and 55 isolates carried the Int2 gene. Int3 genes were absent in all the isolates. Their finding revealed a strong correlation between integrons (especially Int1) and multidrug resistance. Integron-mediated multidrug resistance in extended beta-lactamase-producing isolates of *Klebsiella pneumonia* was investigated by Mobarak-Qamsari et al., [**57**] using 104 clinical isolates. They detected the occurrence of integron classes 1, 2, and 3 in a PCR analysis of the isolates, and found that 50 isolates were related to the production of extended-spectrum beta-lactamases. Out of the 50, 22 carried integron class 1 and 2 and none carried class 3 integron genes. They also found that integron-harbored isolates were significantly associated with higher rates of multi-antibiotic (aztreonam, ceftazidime, cefotaxime, cefepime, kanamycin, tobramycin, norfloxacin, and spectinomycin) resistance in extended-spectrum beta-lactamase-producing *K. pneumonia*. blaTEM-1 beta-lactamase gene is the highly expressed gene in the CBD-resistant *S*. Typhimurium strain among all the gene considered in this study (**Figure 7A and Figure 8**). And this gene (blaTEM-1 beta-lactamase) is one of the regularly documented antibiotic resistance determinants globally. It is known to exhibit stringent resistance to penicillin and cephalosporins [**58**]. Several studies demonstrated that blaTEM-1 beta-lactamase plays a crucial role in antibiotic resistance development in some clinically important pathogens such as *Salmonella, Neisseria gonorrhoeae, E. coli, Klebsiella pneumonia, and Pseudomonas aeroginosa* [**59, 60, 61**]. Yang et al., [**62**] reported that all their isolates carried the blaTEM gene, which was the cause of most of the resistance occurring in *Salmonella* in a large breeder farm in Shandong China when they investigated the occurrence and characterization of *Salmonella* isolates from large-scale breeder farms. Similarly, a study carried out in Thailand to determine the antimicrobial genes in *Salmonella* enterica isolates from poultry and swine indicates that the *blaTEM* gene along with other genes such as *cmlA, tetA, dfrA12, sul3, and aada1* was detected in majority of the strain [**62**]. Two variants of *the blaTEM* gene were also prevalent among ampicillin-resistant *E. coli* and *Salmonella* isolated from food animals in Denmark, which were designated *blaTEM-127* and *blaTEM-128*.

The spy protein is a chaperone, that is expressed and localized in the bacterial periplasm of *E. coli* or *S*. Typhimurium during spheroplast formation, or by exposure to protein denaturing conditions [**63**]. Interestingly, spy induction has been shown to require regulatory proteins such as BaeR and CpxR, which are part of the major envelope stress response systems BaeS/BaeR and CpxA/CpxR. Thus, scrutinizing spy transcription could be a good determinant of the mode of activity of an antimicrobial agent, particularly, if it can cause envelope disruption in *E. coli* or in *S*. Typhimurium [**64**].

In this work, we demonstrated for the first time that CBD-resistant strain of *S*. Typhimurium showed higher spy expression (**Figure 7C**), thus, CBD could be used to induce spy transcription in *S*. Typhimurium. It has been previously demonstrated that agents that induced envelope stresses, including ethanol, polymyxin B, and spheroplasting, cause upregulation in the expression of the spy gene in both *S*. Typhimurium and *E*. coli [63, 64, **65**]. In this study, the spy gene which is known to be expressed mainly in the periplasmic space under envelope stress conditions, showed significantly higher expression in the resistant strain of *S*. Typhimurium which has been exposed to CBD than the susceptible strain which did not receive CBD treatment. Thus this work corroborates with other works that have shown that agents that causes envelope stress activates the transcription of spy, which then acts as envelope stress sensor. While previous work in our laboratory demonstrated that CBD caused envelope disruption [**28, 31**], in this work, it is plausible to infer that CBD-resistant strain of *S*. Typhimurium which showed higher spy expression were due to the induction of the spy gene by CBD. It has been demonstrated that the spy gene could be used as a whole-cell biosensor for testing antimicrobials that target bacterial cell envelope (**66, 67, 68**). The spy gene interact with other closely related genes such as cpxA, cpxR, mdtA, dsbA, ybaJ, nlpB, and STM2535 (**Figure 9B**). The nlpB are nonessential genes that encodes lipoproteins and plays crucial role in cell envelop interigity as was elucidated by Onufryk et al, [**69**]. Jing et al [**70**] analyzed the role of *cpxA* mutations in *Salmonella enterica serovar* Typhimurium resistance to aminoglycosides and β-Lactams using site-directed mutants and an internal in-frame mutants and found that cpxA and cpxR deletion mutants have opposing modulation of resistance to AGAs and β-lactams. Thus, their findings reveal that all cpxA mutations increases resistance to AGAs and β-lactams significantly. From this study, it can be safe to infer that these genes together with the spy gene plays a significant role in *S*. Typhimurium resistance to CBD.

Sterols are an essential part of lipids of the membranes, and sterols in high concentrations are thought to have beneficial biophysical roles on the membrane. In this study we analyzed the abundance of three different kinds of sterols (Ergosterols, myristic, palmitic, palmitoleic, steric, erucic and oleic acids) in both the CBD-resistant and CBD-susceptible strain of *S*. Typhimurium (**Figure 1B, C D**). Sterols are often one of the most abundant biological membranes of eukaryotes and mammals. This study shows higher abundance of myristic and palmitic acid, Ergosterols and oleic acids in the CBD-resistant *S*. Typhimurium strain as compared to the CBD-susceptible *S*. Typhimurium strain. The abundance of membrane sterols signifies the ability of the *S*. Typhimurium to resist CBD. This is agreeing with findings by Tintino et al [**71**] who demonstrated that cholesterol and ergosterols influences the activity of *Staphylococcus aureus* antibiotic efflux pump.

## 5. Conclusions

In summary, for the first time, we demonstrated that CBD-resistance development in *S*. Typhimurium might be caused by several structural and genetic factors. These structural factors were demonstrated here to include LPS and cell membrane sterols, which showed significant differences in abundances on the bacterial cell surfaces between the CBD-resistant and CBD-susceptible strain of *S*. Typhimurium. Specific key genetic elements implicated for the resistance development investigated included fimA, fimZ, int2, ompC, blaTEM, DNA recombinase (STM0716), leucine-responsive transcriptional regulator (lrp/STM0959), and the spy gene of *S*. Typhimurium. In this study, we revealed that blaTEM might be the highest contributor to CBD-resistance, indicating the potential gene to target in developing agents against CBD-resistant strain of *S*. Typhimurium. To fully understand the mechanisms of resistance development, it will be crucial to investigate all possible genes that might play role in this event, thus, we are currently employing a follow up study using throughput RNA sequencing to unravel all the genetic factors blamable in *S*. Typhimurium development of resistance against CBD.

## Author Contributions

JAA: conceptualization. II and JAA: methodology, software, visualization, and writing–original draft preparation. JAA, OA, and DAA: validation and resources. JAA, JX, DAA, RKB, and OA: formal analysis. II, JAA, DAA, JX, RKB, and OA: investigation. JAA, JX, and OA: data curation. II, JAA, JX, DAA and OA: writing–review and editing. JAA and OA: supervision. OA: project administration and funding acquisition. All authors contributed to the article and approved the submitted version.

## Data Availability Statement

The original contributions presented in the study are included in the article; further inquiries can be directed to the corresponding authors.

## Acknowledgments

We acknowledge Alabama State University, C-STEM for supplies and Laboratory space. The authors acknowledge receiving funding from the United States Department of Education, Title III-HBGI-RES.

## Disclaimer/Publisher’s Note

All claims expressed in this article are solely those of the authors and do not necessarily represent those of their affiliated organizations, or those of the publisher, the editors, and, the reviewers. Any product that may be evaluated in this article, or claim that may be made by its manufacturer, is not guaranteed or endorsed by the publisher.

